# Longitudinal study of persistence in professional development outcomes of early career biology faculty

**DOI:** 10.1101/785857

**Authors:** Nathan C Emery, Jessica Middlemis Maher, Diane Ebert-May

## Abstract

The diversity of teaching professional development (PD) programs that occurred over the last few decades merits our collective attention to assess the impact of these programs over time. In general, the goal of PD programs is that participants continue to practice what they learn in the long term. However, we do not know the degree to which the outcomes of these programs were achieved and ultimately persist. We tracked postdoc participants from the Faculty Institutes for Reformed Science Teaching (FIRST) IV program into their current position as early-career faculty at institutions across the United States. We assessed their teaching approaches, practices, and student perceptions of the learning environment over 6-10 years. Additionally, the FIRST IV faculty were paired with colleagues of similar status in the same departments. We found that professional development outcomes from the FIRST IV program persisted over time and across a significant career transition, from postdoc to faculty. These participants not only maintained their student-centered practices, but were significantly more student-focused than their peers. Lastly, we found that faculty approaches to teaching were correlated with observed teaching practices in the classroom for both groups of faculty. These results provide compelling evidence for the success of the FIRST IV program and the long-term persistence of professional development outcomes.

## Introduction

Teaching professional development (PD) programs have proliferated during the last few decades, with the goal of facilitating measurable change in higher education. However, we know little about the long-term outcomes of faculty professional development efforts (Stes et al. 2009, Steinert et al. 2016, Manduca 2017, Connolly et al. 2018). Given the significant investment in time and resources that these PD programs represent, it is critical that we assess the longitudinal impact on instructional practices and additional program outcomes.

Many teaching PD programs focus on convincing instructors to change the way they teach (Hilborn 2013, Brownell and Tanner 2012), by emphasizing evidence-based practices that support student learning through collaborative and inquiry-based learning. Often, achieving this goal requires shifting faculty away from teacher-centered approaches and towards learner-centered teaching, whereby students engage in their own knowledge construction and focus on conceptual learning (e.g. Cooper et al. 2015). The expected outcomes of teaching PD are transformed student-centered courses (Ebert-May et al. 2015) and changes in faculty attitudes, approaches, and practices in the classroom (Pfund et al. 2009, Manduca et al. 2017, Connolly et al. 2018). Approaches to teaching revolve around the instructor’s intentions and strategies for teaching a given course (Trigwell and Prosser 2004), while teaching practices are how these strategies manifest in a classroom setting (Sawada et al. 2002).

Evaluating these outcomes is essential to understanding how teaching PD is impacting the larger landscape of higher education (Emery et al. 2019). Several large-scale PD programs have reported changes in faculty soon after program completion (Derting et al. 2016, Pfund et al. 2009, Manduca et al. 2017, Connolly et al. 2018). For example, participants in the Summer Institutes program reported using scientific teaching approaches more frequently after completion of the program (Pfund et al. 2009). Additionally, former participants of the “On the Cutting Edge” Program reported using more active learning exercises and teaching observations revealed greater student engagement in classrooms, suggesting a shift to more learner-centered teaching practices (Manduca et al. 2017).

The Faculty Institutes for Reformed Science Teaching (FIRST) IV program, another national-level PD effort, was implemented from 2009 to 2013 and sought to train biology postdoctoral scholars in learner-centered teaching practices (Ebert-May et al. 2015, Derting et al. 2016). The FIRST IV program combined an intensive 3-4 day institute (during the summer) with collaborative, mentored development and teaching of a transformed course at postdocs’ home institutions (during the academic year). This model was repeated twice for each participant, resulting in two years of iterative PD for each postdoc. Shortly after completion of FIRST IV, participants reported using more student-focused approaches to teaching and were directly observed to use more learner-centered classroom practices compared to their peers (Derting et al. 2016).

A key design feature of FIRST IV was the focus on the development of postdocs as future faculty. If this PD program was to have a lasting impact on undergraduate learning, then FIRST IV participants would need to maintain their learner-centered focus over time, and transfer the approaches and practices that they established as postdocs into their teaching as faculty. Testing this hypothesis, which is critical to the assessment of program impacts, requires a longitudinal approach. Longitudinal studies compare instructors to themselves over a length of time and are valuable to education research because they reveal maintenance of behavior, long term impacts to teaching (White and Arzi 2005), and track changes in behavior (Roberts et al. 2006).

In this study, we define longitudinal as comparing assessments of instructors from professional development as postdoctoral scholars to assessments of instructors as faculty. We are not defining this as a set time period, but rather as a shared set of events over time (postdoc PD experience followed by the career transition to faculty). From the beginning to the end of this study, there was a minimum of three years and a maximum of eight years between completion of the FIRST IV program and this study. Our data collection was timed to capture the whole range of early-career (pre-tenure) faculty years. Few studies have examined faculty instructors in the ‘novice’ or early-career stage (Stes et al. 2007, Connolly et al. 2018), but this is a critical time for the development of teaching practices (Austin 2011).

Most longitudinal studies of faculty teaching have relied on qualitative data and self-reported surveys (Dennick 2003, Knight et al. 2005, Stes et al. 2007, Tennill and Cohen 2013, Stewart 2014) that have inherent drawbacks (Bodner 2016, Porter et al. 2014, Saxon et al. 2003). In this study, we evaluate a combination of faculty self-reported approaches to teaching, observed teaching practices, and student perceptions of the classroom environment among both FIRST IV faculty and a set of comparison faculty within the same academic departments (Emery et al. 2019). The questions we seek to answer are threefold: (1) How are PD outcomes maintained over time, as measured by teaching approaches and practices used by FIRST IV faculty? (2) How do teaching approaches, practices, and student perceptions compare between FIRST IV faculty and paired colleagues in their respective departments? (3) How well do self-reported teaching approaches reflect observed practices in the classroom and student perceptions of the classroom environment? We hypothesized that FIRST IV faculty would continue to use the same degree of learner-centered teaching practices over time, and to a greater extent than their peers. Specifically, we predicted that former FIRST IV participants would report a more student-focused approach to teaching, demonstrate higher student engagement in the classroom, and that these differences would be reflected in student perceptions of the classroom environment.

## Methods

### Study Design

For three consecutive years, this study assessed teaching approaches, teaching practices, and student perceptions of the learning environment by administering survey instruments to faculty participants, video recording classroom teaching practices, and distributing survey instruments to students in participants’ courses (see Emery et al. 2019). In addition, faculty participants completed a background survey at the beginning of the study. All surveys were distributed through the Qualtrics Survey system and video recordings of classrooms were conducted by participants and uploaded to secure cloud servers.

### Participants

This study solicited former participants of the FIRST IV program who, prior to the start of the study, were all early-career faculty at academic institutions across the country. We preferentially contacted tenure-track faculty at different types of institutions (doctoral, masters, baccalaureate, and associate levels). Participants in the longitudinal study came from seventeen doctoral universities, eleven master’s colleges & universities, four baccalaureate colleges, and two associate colleges according to the Carnegie classification system (Carnegie Foundation for the Advancement of Teaching, 2011). The institutions are located in 24 states from all regions of the US including New England, Great Lakes, South, Midwest, Mountain West, and West Coast. Institution size ranged from 800 to 50,000 enrolled students. Each participating FIRST IV faculty was asked to seek out a paired colleague for the study, that is, someone at a similar career stage in their same department, who had not participated in the FIRST IV program. The goal was to establish a comparison group of faculty who were similar to the FIRST IV faculty across dimensions except for their experiences in professional development. This study design allows us to ask questions about how the selected group (FIRST IV faculty) are different from the comparison group (Comparison faculty; Fraenkel et al. 2011). Each participant who collected and submitted data for the three consecutive years of the study was given an honorarium, and 40 FIRST IV faculty and 40 paired comparison faculty completed the study. Some participants did not collect data for all three years and other participants were recruited mid-study. One FIRST IV participant switched institutions mid-study and data collection continued at the new institution with a new paired comparison faculty participant.

### Past data collection for FIRST IV participants

Data were also available from FIRST IV participants when they originally completed the PD program (6 to 10 years ago). Survey and classroom observation data were collected for the courses taught by postdocs while participating in the FIRST IV program. This study aggregated past participant and student-reported data collected using the Approaches to Teaching Inventory (ATI; Trigwell and Prosser, 2004), Reformed Teaching Observation Protocol (RTOP; Sawada et al. 2002), and the Experiences in Teaching and Learning Questionnaire (ETLQ; Entwistle et al. 2002) from Ebert-May et al. (2015). Instrument scores were averaged for each participant. The number of courses taught per participant during FIRST IV varied from one to three courses. These data will be referred to as “past” data, whereas all data from the current study will hereafter be considered “present” data. These two timestamps, which are only available for FIRST IV participants, allow us to make longitudinal comparisons of teaching approaches and practices.

### Background and course information

We administered a background survey through Qualtrics to each faculty participant as soon as they consented to participate in the longitudinal study. This survey collected information pertaining to the academic/training background of each participant, their current academic position, and knowledge, experience, and confidence relating to pedagogical practices and previous teaching activities. We collected course information from faculty at the beginning of each course. This included, for example, course size, student type (major/non-major), and course type (lecture/lab).

### Faculty teaching approach

For each course taught, a faculty participant completed the ATI survey on Qualtrics at the beginning of their course. This instrument assesses the extent to which an instructor uses teacher-focused and student-focused teaching approaches. It is a self-reporting instrument and is useful for assessing instructor approach for courses in different disciplines (Williams et al. 2015). The ATI produces two subscale scores, Conceptual Change/Student-Focused (CCSF) and Information Transfer/Teacher-Focused (ITTF). Instructors who teach with a primarily CCSF approach focus on engaging students with deep approaches to build and modify their conceptual understanding of the course material. Instructors who teach with an ITTF approach transfer information to their students and focus on competency (Trigwell & Prosser 2004). The two subscales are independent of one another; thus two scores were calculated per participant per course.

### Observations of teaching practices

For each course, we asked study participants to collect video recordings of two class sessions. We advised participants to allow at least one week between each video recording and to focus the recorder on the slide presentation and them as they taught the class and interacted with students. Audio was particularly important to capture on the recording. In some instances, participants were only able to record one video per course. All recordings were de-identified before review. As in the previous study of FIRST IV participants (Ebert-May et al. 2015), the Reformed Teaching Observation Protocol (RTOP; Sawada et al. 2002) was used to rate teaching videos on the extent to which the course was student-centered and transformed. RTOP is a teaching observation tool designed to assess teaching reform through use of learner-centered practices (Sawada et al. 2002). The tool suggests using at least two observations per participant, although recent work suggests that at least four observations are necessary for a reliable characterization of teaching (Stains et al. 2018). For this study, there were two observations per course each year for each participant, resulting in a maximum of six observations per participant over the course of the study.

To avoid potential bias due to visual recognition of instructors, we recruited independent video raters with expertise as doctoral-trained biologists from several academic institutions. Raters were trained in the use of the RTOP instrument until they calibrated with the trainer and the rater cohort. Over the course of the study, 12 raters were involved in reviewing videos, and these raters were calibrated to one another with an intra-class correlation coefficient (ICC; Gwet 2010) of 0.77 over six videos. We randomly assigned each de-identified study video to two raters, and the scores were averaged. Any video scores with a standard deviation above seven were assigned to a third-party rater, and the subsequent two closest scores were used in the analyses.

### Student perceptions of the classroom and scientific teaching practices

For each course, faculty participants administered the ETLQ to their students near the end of the term. Students completed the questionnaire on Qualtrics and submitted their responses through a faculty participant-specific link. Students entered their student ID so that faculty could receive a list of survey completion, but not their responses. Some faculty chose to incentivize students with course credit or extra credit for the survey while others chose not to assign points to the survey. We chose to analyze several of the subscales found in the ETLQ including: Deep Approach, Surface Approach, Alignment, Choice, Encouraging High Quality Learning, Organization, Structure and Content, and Peer Support. Previous studies found the ETLQ to be robust enough to pull out a subset of scales and slightly modify the questionnaire (Mogashana et al. 2012, Stes et al. 2013, Nigh et al. 2015). The subscales in our study are aligned with the research questions and reflect student perceptions of a student-focused classroom. For example, a Deep Approach to teaching is perceived by students as relating ideas and using evidence, while a Surface Approach consists of memorizing without understanding and compiling fragmented knowledge. Alignment is how well the concepts and practices that students were taught matched the goals of the course. Choice refers to students’ perceptions about choices in their own learning. Encouraging High Quality Learning is whether students felt they were prompted in class to self-reflect and learn about how knowledge is developed. Organization, Structure and Content is the perceived clarity of course objectives, organization, and flow of the classroom. Peer Support is how students supported each other and felt comfortable working with others (Entwistle et al. 2002).

In addition to student perceptions of the classroom environment, we assessed student perceptions of the frequency of scientific teaching practices they experienced in the classroom using the Measurement Instrument for Scientific Teaching (MIST; Durham et al. 2017). This instrument was published after the first year of data collection and thus was distributed in courses during year two and three of this study. Faculty participants distributed the MIST to students along with the ETLQ at the end of each course.

All student survey data were filtered to remove incomplete responses, duplicate responses, and poor responses from students (e.g. entering 5’s for each item). The response rate ranged from 7.6% to 100% with an average response rate of 67.6% across all three years of the study. We examined the variance in student responses to choose a percent response rate threshold. For five courses with large enrollments and high percent response rate (>90%), subscale scores were resampled at 10, 20, 30, 40, and 50% response rates. The subsequent standard deviations in student responses were plotted by resampling and it was determined that a 30% response rate was the lowest threshold indistinguishable from 50%, an accepted threshold of response in the literature (Fosnacht et al. 2017). Thus, ETLQ and MIST data were only analyzed from courses with >30% response rate. This resulted in the removal of 26 courses (out of 212) from the present ETLQ and MIST data analysis.

### Analysis

Past (from FIRST IV) and present (all current courses) scores for ATI, RTOP, and ETLQ scores were averaged per participant to encompass their teaching approach, practices, and student perceptions. We tested all past and present variables for normality prior to statistical analysis. Only the ITTF subscale of the ATI had a non-normal distribution. For the ATI and ETLQ subscales, which are ordinal, we ran a non-parametric paired Wilcoxon sign-rank test between the past mean scores and the present mean scores for FIRST IV Faculty. For RTOP scores we ran a paired Student’s t-test between the past scores and the present scores.

To determine the difference in ATI and ETLQ subscales between the FIRST IV faculty and the comparison faculty we ran paired Wilcoxon sign-rank tests using the within-department pairings of FIRST IV faculty and comparison faculty. We similarly used a paired Student’s t-test to compare RTOP scores between the two groups.

It is possible that ATI, RTOP, and ETLQ scores have shifted over the three years of data collection (2016-2019). To determine if scores changed significantly during the study, we took instrument subscale scores from each course and ran simple linear regressions for all instrument scores over the three years of the study. We ran separate regressions for FIRST IV faculty and comparison faculty. Additionally, it is possible that time as a faculty instructor or years of teaching experience may influence their approach and teaching practice. We analyzed the relationship for self-reported “years as faculty” and “years of teaching experience” with ATI subscales and RTOP results. Linear regressions estimated how “years as faculty” or “years of teaching experience” were related to teaching approach and teaching practice.

Instructor approaches and teaching practices in the classroom are also possibly related to one another. We took the mean score per participant (across all years of participation) and ran a simple linear regression of both ATI subscales (CCSF and ITTF) and RTOP to determine how approaches may be reflected in teacher practices in the classroom. Additionally, we regressed RTOP scores with ETLQ and MIST subscale scores to determine if teacher practices were detectable in self-reported student perceptions of the classroom.

All analyses were examined for differences in responses according to self-reported instructor gender identity.

## Result

### Description of participants and courses

We first verified that FIRST IV faculty and comparison faculty were similar in relation to demographics, academic appointment and teaching experiences. The majority of both groups of faculty identified as women (60% FIRST IV; 70% comparison). The percent of pre-tenure faculty in both groups was the same (86%) and both had approximately six years of teaching experience. Participation in prior professional development (PD) activities was significantly higher for FIRST IV faculty than for comparison faculty, with more than twice the number of hours reported (Table 1; ANOVA, p-value < 0.001).

**Table 1.** Instructor identity, academic appointment and teaching experience at the beginning of the study.

| Instructor descriptors | FIRST IV (40) | Comparison (40) |
| --- | --- | --- |
| Identify as Woman / Man | 24 / 16 | 28 / 12 |
| Tenured / Tenure track / Non-tenure track | 5 / 32 / 3 | 5 / 31 / 4 |
| Percent appointment: Teaching (mean ± SD) | 53.2 ± 19.6 | 53.1 ± 20.4 |
| Percent appointment: Research (mean ± SD) | 34.9 ± 20.4 | 31.6 ± 19 |
| Years as faculty prior to study (mean ± SD) | 3.4 ± 1.6 | 2.9 ± 3.1 |
| Years of teaching experience prior to study (mean ± SD) | 6.5 ± 3.4 | 6.4 ± 6.1 |
| Highest engagement PD activity in hours (mean ± SD) | 90.44 ± 58.8 | 39.45 ± 37.4 |

Course characteristics were also similar across the two groups of faculty (Table 2). Course enrollment ranged from 3 to 624 with a mean and standard deviation for FIRST IV faculty of 72 ± 106 and for Comparison faculty 61 ± 62. Most courses in this study were designed for undergraduate science majors and the distribution of courses for majors vs. non-majors was similar between the FIRST IV faculty and Comparison faculty.

**Table 2.** Course and student information for 2016-2019.

| **Course Category** | **FIRST IV (N=111)** | **Comparison (N=105)** |
| --- | --- | --- |
| Large lecture (>60 students) | 18 | 16 |
| Large lecture with lab | 11 | 13 |
| Small lecture (<60 students) | 28 | 25 |
| Small lecture with lab | 49 | 44 |
| Small discussion course | 1 | 4 |
| Other course type | 4 | 3 |
| Course enrollment (mean ± SD) | 70.8 ± 105 | 61.6 ± 61.3 |
| **Type of students** |  |  |
| Majors | 70 | 70 |
| Non-Majors | 12 | 7 |
| Both Majors and Non-Majors | 29 | 28 |

Subscale scores for teaching approach (ATI; Trigwell and Prosser 2004), teaching practice (RTOP; Sawada et al. 2002), and student perceptions of the learning environment (ETLQ; Entwistle et al. 2002) did not significantly change over the three years of the study for either the FIRST IV faculty or the Comparison faculty. There was no significant relationship between “year of study” and instrument scores. Additionally, there was no significant relationship between “years as faculty” or “years of teaching experience” and ATI subscale or RTOP scores.

### Persistence of teaching practices (past and present)

FIRST IV faculty reported similar teaching approaches (ATI) in courses at the end of the FIRST IV professional development program (2011 or 2013) and courses in this study (2016-19) (Figure 1A). No significant differences in FIRST IV faculty’s support for the use of conceptual-change/student-focused (CCSF) or information-transmission/teacher-focused (ITTF) teaching strategies in their courses were noted between past and present values of the ATI subscales: CCSF (paired Wilcoxon sign-test, p-value = 0.51) or ITTF (paired Wilcoxon sign-test, p-value = 0.103). This is evidence of persistence in faculty teaching approach. For both past and present ATI scores, the CCSF subscale score was significantly higher than the ITTF score (*past*: paired Wilcoxon sign-test, p-value < 0.001; *present*: paired Wilcoxon sign-test, p-value <0.001). There was no effect of gender on persistence of teaching approach.

**Figure 1.**
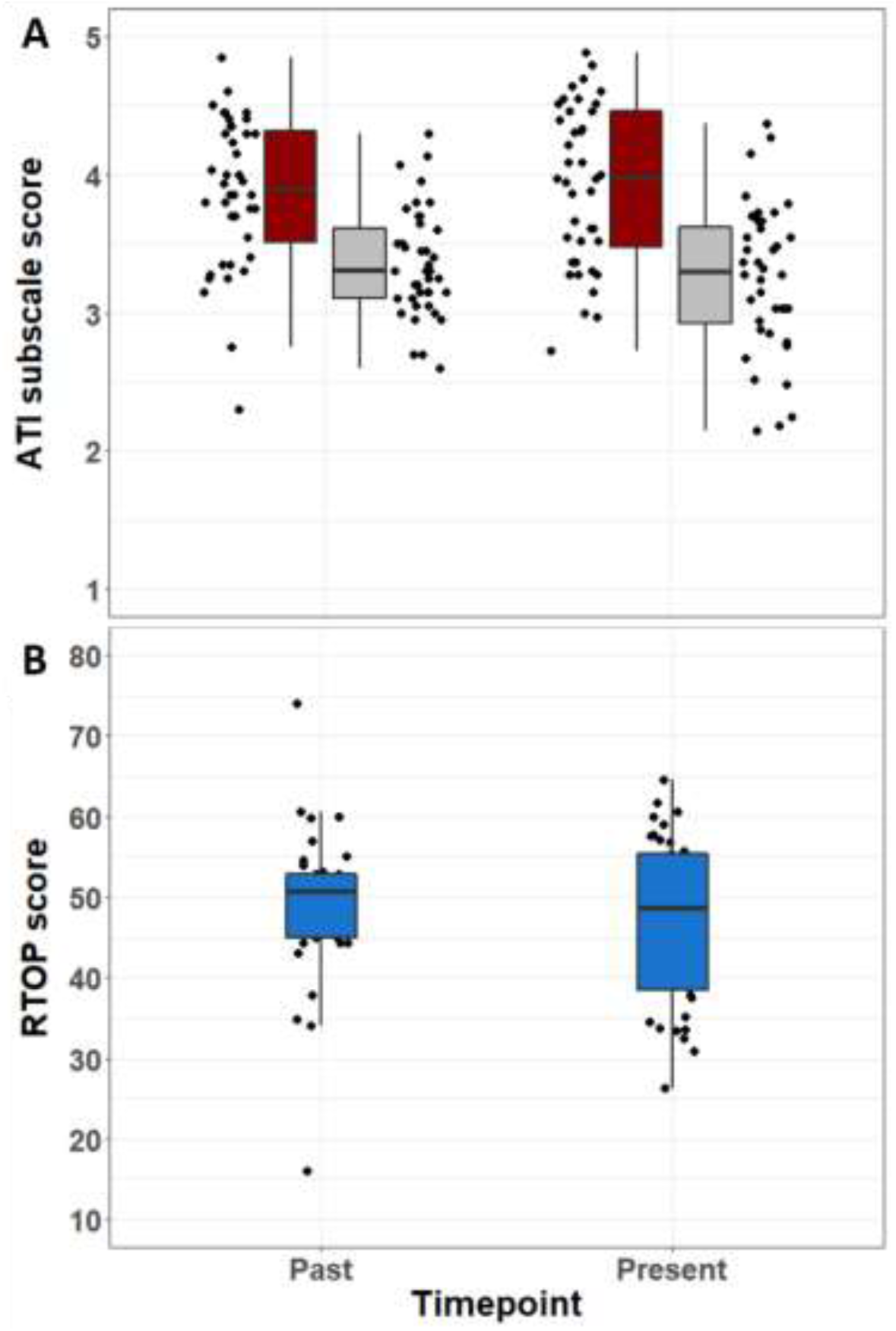
Past and present teaching approach and teaching practice. [A] Past ATI scores compared to present scores for FIRST IV study participants. Each point represents the mean ATI score for Conceptual Change/Student-Focused (CCSF; Red) and Information Transfer/Teacher-Focused (ITTF; Gray) subscales per participant. The boxes represent the interquartile range. [B] Past RTOP scores compared to present scores for FIRST IV study participants. Each point represents the mean RTOP score per participant. The boxes represent the interquartile range.

The mean Reformed Teaching Observation Protocol (RTOP; Sawada et al. 2002) score of videos for FIRST IV faculty (47.5 ± 11.5) was within RTOP category 3, which is characterized by significant student-engagement with some minds-on as well as active involvement nearly half of the class period. RTOP scores were not significantly different between the present study and past results (paired Student’s t-test, p-value = 0.326). Faculty demonstrated similar teaching practices in the present as they had in the past (Figure 1B), evidence of persistence in observed teaching practices post-professional development. There was no effect of gender on persistence of teaching practice.

Despite teaching at different institutions with different student populations, there were no significant differences between past and present ETLQ subscale scores for FIRST IV faculty participants (paired Wilcoxon sign-tests). All differences in subscales were non-significant, including Deep Approach to learning and studying (p-value = 0.107), Surface Approach to learning and studying (p-value = 1), Alignment of teaching and learning (p-value =0.066), Choice about learning (p-value = 0.75), Encouraging High Quality Learning (p-value = 0.058), Organization, Structure and Content (p-value = 0.063), and Peer Support (p-value =0.468). The only gender difference observed in ETLQ subscales was in Encouraging High Quality Learning. Students of female instructors reported significantly higher values in the present study than in the past (p-value = 0.038); whereas students of male instructors reported no significant differences in Encouraging High Quality Learning (p-value = 0.97).

### Teaching approaches and practices during present study

FIRST IV faculty and Comparison faculty did not significantly change their approaches to teaching and teaching practices over the three years of this study. ATI subscale scores did not significantly change over three years for CCSF (linear regression, FIRST IV faculty, r^2^ = 0.004, p-value = 0.517; Comparison faculty, r^2^ = 0.005, p-value = 0.219) or ITTF (linear regression, FIRST IV faculty, r^2^ = 0.001, p-value = 0.713; Comparison faculty, r^2^ = 0.005, p-value = 0.471). Also, RTOP scores did not significantly shift over the course of this study (linear regression, FIRST IV faculty, r^2^ < 0.001, p-value = 0.965; Comparison faculty, r^2^ = 0.004, p-value = 0.554). Additionally, there was no significant relationship between RTOP scores and years as a faculty member (linear regression; r^2^ = 0.002, p-value = 0.68) or years of teaching experience (linear regression; r^2^ = 0.001, p-value = 0.756).

### Faculty comparisons

FIRST IV faculty had a greater student-focused approach in their courses and a lower teacher-focused approach than the Comparison faculty (Table 3). FIRST IV faculty had significantly higher CCSF scores than Comparison faculty (paired Wilcoxon sign-test, p-value = 0.039) and lower ITTF scores (paired Wilcoxon sign-test, p-value = 0.016). For Comparison faculty, there was no significant difference between the CCSF and ITTF subscales (Wilcoxon sign-test, p-value = 0.07).

**Table 3.** Mean and standard deviation of subscale scores from different assessment types for FIRST IV faculty and Comparison faculty. * indicates statistically significant differences between faculty groups for p <0.05 (ATI, paired Wilcoxon signed rank test; RTOP, MIST, paired Student’s t-test).

| **Assessment Type** | **FIRST IV Faculty** | **Comparison Faculty** |
| --- | --- | --- |
| ATI: Conceptual Change/Student-Focused (mean ± SD) | 3.98 ± 0.6* | 3.76 ± 0.64 |
| ATI: Information Transfer/Teacher-Focused (mean ± SD) | 3.21 ± 0.56* | 3.51 ± 0.54 |
| Reformed Teaching Observation Protocol (mean ± SD) | 48.03 ± 11.67* | 38.62 ± 9.67 |
| MIST: Composite Score (mean ± SD) | 58.4 ± 8* | 53.2 ± 9.4 |

FIRST IV faculty had significantly higher RTOP scores than Comparison faculty (Table 3, paired Student’s t-test, p-value < 0.001). There were also more FIRST IV faculty in RTOP categories 3 (previously defined) and 4 (active student participation and critique of science processes) than Comparison faculty (Figure 2). These categories indicate significant student involvement in open-ended inquiry and scientific practices in the classroom.

**Figure 2.**
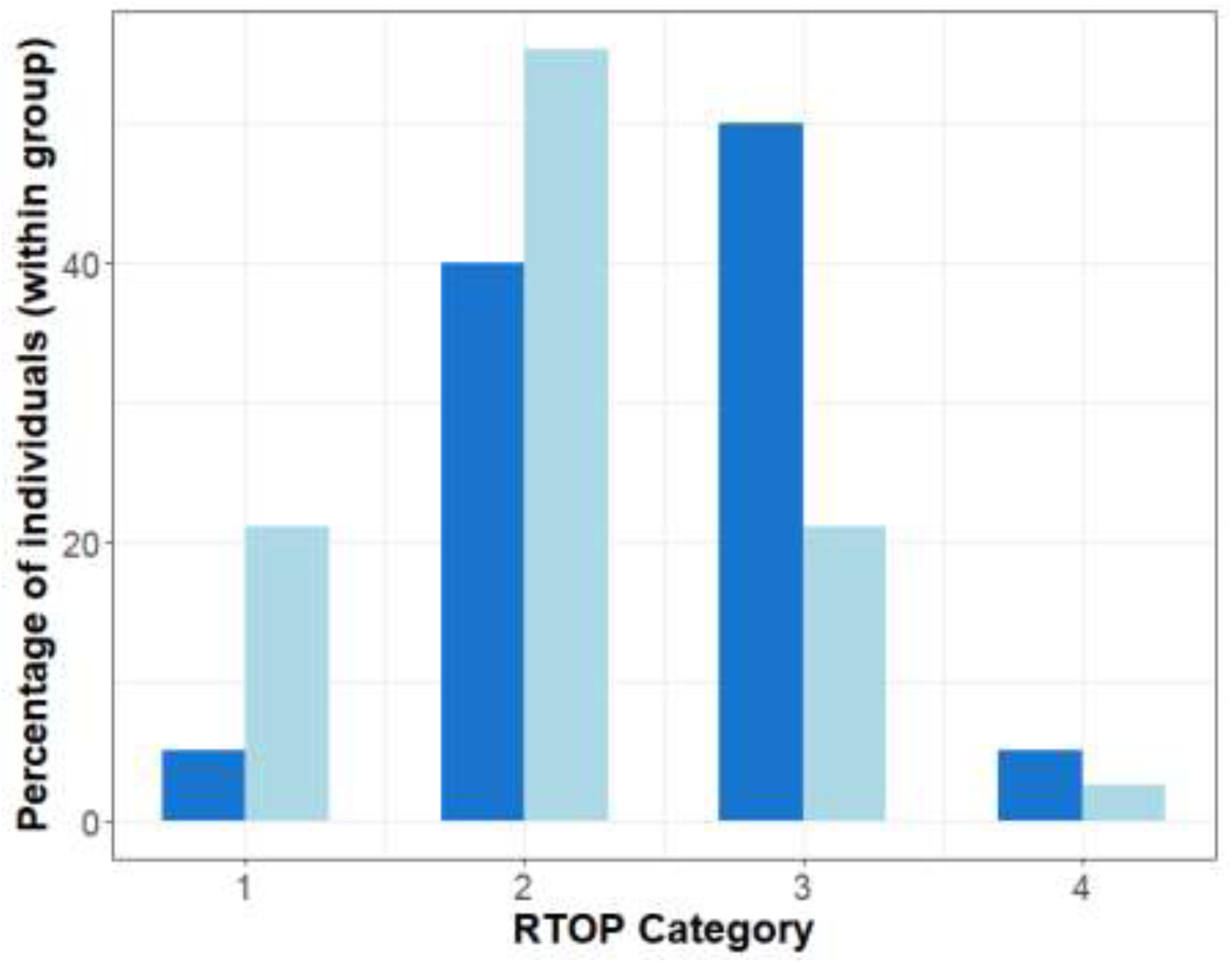
Comparison of RTOP categories for FIRST IV faculty (Dark Blue) and Comparison faculty (Light Blue). Each bar represents the percent of faculty (within a group) whose mean RTOP score fits within a particular category.

Student perceptions of the classroom did not significantly differ between the two groups of faculty for all of the ETLQ subscale scores. Despite no differences in the ETLQ, students did report a greater frequency of scientific teaching in the classroom for FIRST IV faculty than Comparison faculty according to the Measurement Instrument for Scientific Teaching (MIST) composite score (paired Wilcoxon sign-test, p-value = 0.034).

### Relationship between reported teaching approaches and observed teaching practices

For both FIRST IV and Comparison faculty, CCSF subscale scores were positively correlated with RTOP scores (Figure 3, FIRST IV: linear regression, r^2^= 0.186, p-value < 0.001; Comparison: linear regression, r^2^= 0.262, p-value < 0.001). Additionally, ITTF scores were negatively correlated with RTOP scores for both FIRST IV (Figure 3, linear regression, r^2^= 0.086, p-value = 0.0014) and Comparison faculty (linear regression, r^2^= 0.176, p-value < 0.001). The relationships between teaching approach and teaching practice were the same for both men and women faculty in the study.

**Figure 3.**
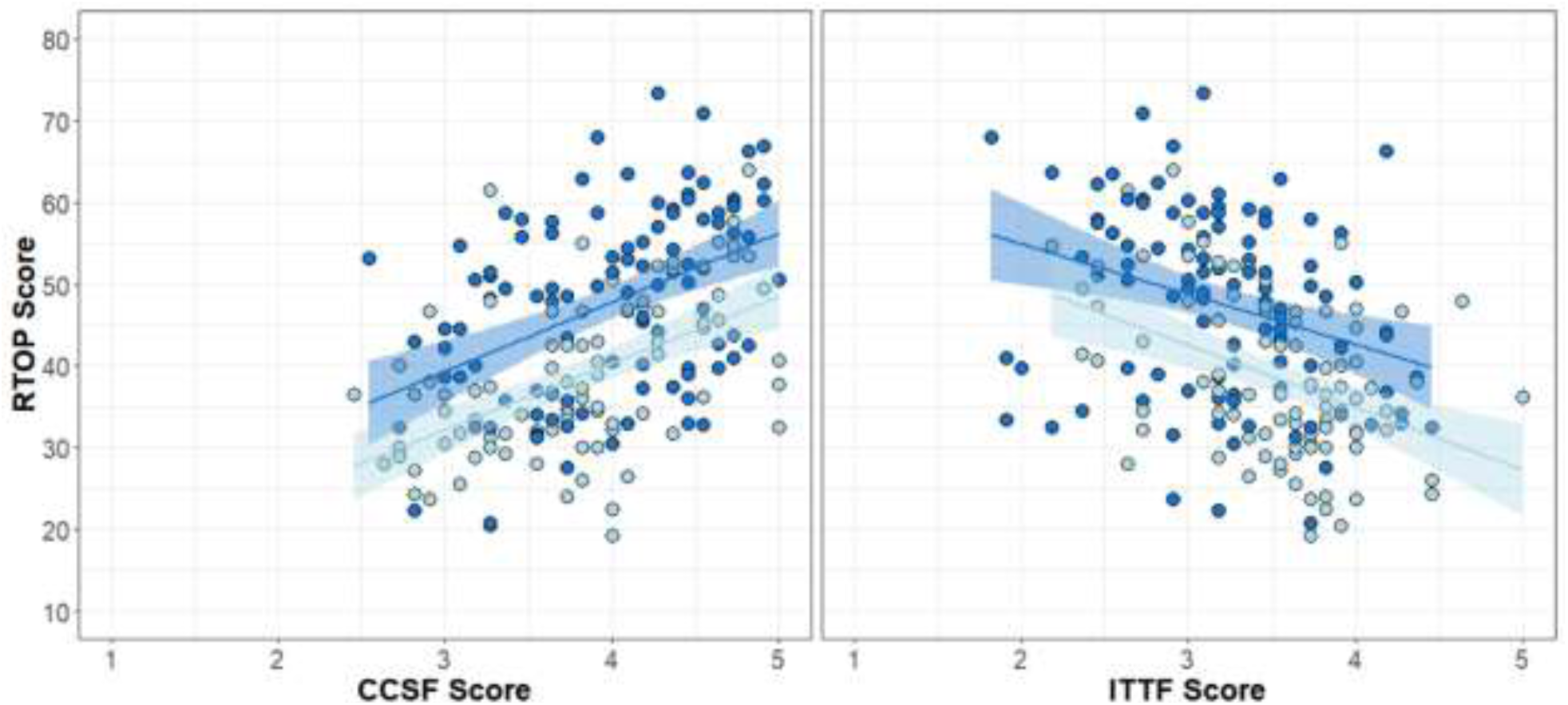
Relationship between ATI subscales Conceptual Change/Student-Focused and Information Transfer/Teacher-Focused and RTOP scores for each course assessed in the study. Each point represents data from a single course for FIRST IV faculty (Dark Blue) and Comparison faculty (Light Blue). The lines represent linear regressions with confidence intervals.

### Alignment of observed teaching practice with student perceptions of the classroom

Across both groups of faculty, there were few relationships between observed teaching practice (measured by RTOP) and student perceptions of the classroom environment as measured by the ETLQ. A negative correlation existed between the RTOP score and the ETLQ subscale Surface Approach (linear regression, r^2^= 0.031, p-value = 0.009), and a positive correlation existed between the RTOP score and the ETLQ subscale Choice (linear regression, r^2^= 0.055, p-value < 0.001). There was a significant positive relationship between student perceptions of scientific teaching practices as measured by the MIST and observed teaching practices measured by RTOP (Figure 4; linear regression, r^2^= 0.194, p-value < 0.001). The relationships between teaching practice and student perceptions of the classroom were consistent across genders. There was no significant gender difference in student survey responses for any of the subscales except for “MIST: Course and Self-Reflection” which had higher scores for male instructors (Student’s t-test, p-value = 0.038) and “ETLQ: Surface approach” which had higher scores for male instructors (Wilcoxon sign-test, p-value = 0.012).

**Figure 4.**
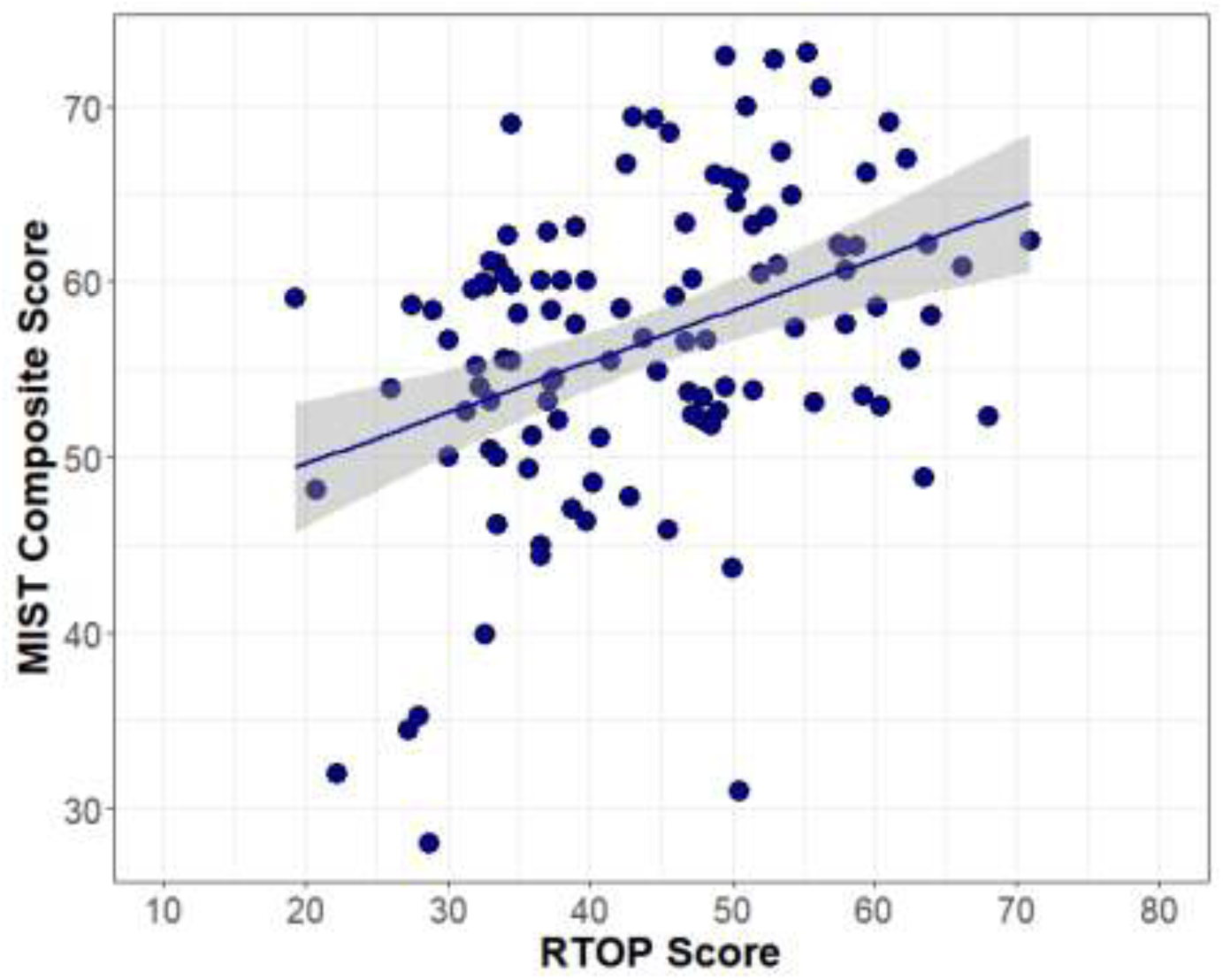
Relationship between observed teaching practices (RTOP) and student perceptions of scientific practices (MIST Composite) for all faculty. Each data point represents a single course with a greater than 30% student response rate for the MIST survey instrument. The line represents a linear regression of the relationship with confidence intervals.

## Discussion

Teaching PD has developed in various formats and scales from broad-reaching programs at the national level, such as FIRST IV (Derting et al. 2016), Summer Institutes (Pfund et al. 2009), Center for the Integration of Research, Teaching, and Learning (CIRTL) (Austin et al. 2008), and On the Cutting Edge (Manduca et al. 2017), to local programs focused on individuals and departments within an institution. A shared goal across all of these programs is to change the way that science is taught at the college level, which requires that participants will continue to practice what they learn in the program.

From 2009-2013, FIRST IV postdoctoral scholars participated in a rigorous teaching development program aimed at cultivating a student-focused approach to teaching that emphasized student engagement with scientific practices to learn concepts/content (Ebert-May et al. 2015). Soon after, many FIRST IV participants were found to have significantly greater student-focused classrooms than their peers (Derting et al. 2016). Results from the present study extend this finding substantially, and suggest that the outcomes of teaching PD can persist across a career transition and well into a faculty instructor’s career in higher education (Figures 1A & B). Additionally, the former PD participants studied here demonstrate significantly greater student-focused approaches and practices in the classroom than their colleagues (Table 3). This study provides powerful evidence of the impact of the FIRST IV model on future faculty teaching development. Results from this study supported our hypothesis that FIRST IV faculty teaching would not significantly change between the PD program and present time, and that FIRST IV faculty would have a more student-focused classroom than matched comparison faculty.

It is possible that faculty teaching practices become more student-focused with more experience, however, our results do not suggest that this is the case for either the FIRST IV faculty or Comparison faculty. Faculty with more teaching experience were not using a more student-focused approach or implementing student-centered practices in the classroom. This suggests that senior faculty may also benefit from professional development (Huston & Weaver 2008). The results of our study echo the call for more longitudinal studies in higher education (White and Arzi 2005, Stes et al. 2009, Henderson et al. 2011, Steinert et al. 2016). By tracking professional development participants over time, we gain a greater understanding of program outcomes and effectiveness of the program itself.

### Persistence of Teaching Development Outcomes

A critical gap in our understanding of teaching development programs are the long-term outcomes for faculty (Stes et al. 2009, Brownell and Tanner 2012, Steinert et al. 2016). Our longitudinal study captured the change in teaching for faculty between three and eight years after completing the FIRST IV program. Faculty participants showed no significant change in their approach to teaching or use of teaching practices (Figure 1A & B). PD outcomes can change for many reasons, including lack of repetition (Henderson et al. 2011), unsupportive communities of practice (Brownell & Tanner 2012, Middleton et al. 2015), and changes to incentives and pressures (Austin 1990). Early career faculty are particularly faced with many new pressures that take time and resources (Austin 2002), potentially causing faculty to reallocate their focus from student-centered teaching practices. Previous longitudinal studies have examined participant outcomes with surveys and interviews and found evidence of persistent PD outcomes (Stes et al. 2007, Tennill & Cohen 2013, Stewart 2014). Our results reinforce past work and confirm the long-term impacts of PD programs with validated survey instruments and teacher observations. In addition, our results suggest that the outcomes from professional development were maintained over a long period of time and through a significant career transition despite the added pressures and barriers to student-centered pedagogy for early-career faculty.

### Trained faculty are more student-focused

A core objective of teaching PD programs is to change instructor attitudes, approaches, and teaching practices. By comparing FIRST IV faculty to paired faculty from the same departments, we were able to assess differences in teaching between FIRST IV faculty and their peers and control for the impacts of departmental culture. There is inherent value in comparing program participants to similar instructors who did not experience the same professional development. The differences observed between FIRST IV and Comparison faculty provide strong evidence of an “effect” of the FIRST IV program. Former participants of the FIRST IV program approached their courses from a more student-focused perspective (Table 3) and implemented more student-centered practices in the classroom (Figure 2). While this finding might be expected for a successful PD program, it is notable given the length of time that has passed since the PD program ended in addition to a significant career transition from postdoc to faculty (typically at a different academic institution).

### Teaching approach translates into teaching practice

There is often a disconnect between what an instructor says they do in a classroom and what they actually do in the classroom (Fung & Chow 2002, Ebert-May et al. 2011). The connection between attitude/beliefs and behavior is not limited to teaching and has been explored thoroughly in behavioral theory (Azjen 1991). Despite observed disconnects in the literature, an instructor’s teaching approach, in theory, influences their teaching practice. Our study results provide evidence of this relationship (Figure 3). Both FIRST IV faculty and Comparison faculty showed a relationship between their self-reported teaching approach (measured by the ATI; Trigwell and Prosser, 2004) and their teaching practices in the classroom as assessed by independent observations (RTOP; Sawada et al. 2002). Student-focused approaches to teaching a course resulted in a more student-centered classroom. Inversely, teacher-focused approaches led to less student-centered classrooms. This suggests that the self-reported teaching approach was reflected in the teaching practices of both faculty groups, despite differences in professional development.

### Student perceptions reflect teaching practices

Students perceptions of learner-centered teaching practices can vary (Machemer and Crawford 2007, Cavanaugh 2011, Durham et al. 2018). The Measurement Instrument for Scientific Teaching confirms results from the ATI and RTOP, that FIRST IV faculty are teaching more student-centered courses than Comparison faculty (Table 3). The MIST instrument was designed to gauge the frequency of scientific teaching practices in the classroom (Durham et al. 2017). Teacher observation scores (RTOP) were positively correlated with MIST composite scores across both groups of faculty (Figure 4). A recent study also found high agreement between instructor and student perceptions of scientific teaching (Durham et al. 2018). Our findings suggest that MIST may be an informative proxy for gauging the learner-centeredness of a course.

The Experiences of Teaching and Learning Questionnaire is designed to assess student perceptions of the teaching-learning environment and how learning occurs in the classroom (Entwistle et al. 2002). It is realistic to hypothesize that these perceptions might change over time and with different student populations. However, there were no significant changes for FIRST IV faculty in any of the ETLQ subscales despite the completely new teaching-learning environment. This could be due to conserved faculty teaching approaches and practices (Figure 1) and/or the ETLQ instrument does not align with RTOP outcomes. Both are possible explanations as there were few relationships observed between the RTOP and ETLQ subscales and few ETLQ subscales differed between FIRST IV faculty and Comparison faculty. Overall, it is unclear how the ETLQ instrument informs our research questions of the learner-centeredness of classrooms.

Despite past findings that students perceive a classroom differently based on the gender of the instructor (MacNell et al. 2015), we found inconclusive results of instructor gender on ETLQ subscales or MIST scores. The ALS: Surface Approach subscale from the ETLQ was slightly higher for male instructors than female instructors. This suggests that men may be teaching their courses from a surface approach, with more memorizing, unreflective studying, and fragmented knowledge. This is slightly contradicted by the MIST: Course and Self-Reflection subscale, whereby male instructors had higher student-reported values. It is unclear if these gender differences are educationally significant given the contradiction and lack of relationships/differences among the other student survey subscales.

## Limitations

While it is likely that the FIRST IV program significantly shifted participant beliefs, attitudes, and practices, it is still unknown how student-focused the participants were prior to professional development. Past data were collected during and towards the end of the FIRST IV program and thus reflect “post” PD outcomes. There is also potentially a self-selection bias from participants who chose to enter the FIRST IV program. The postdoctoral scholars may have been more receptive to evidence-based teaching techniques and/or predisposed to student-focused approaches to teaching prior to the start of FIRST IV. Thus the finding of a lack of change may be indicative of previously-held teaching beliefs or teaching practices established prior to the FIRST IV program.

## Conclusion

Persistent, long-term change is critical to the success of teaching professional development programs. The FIRST IV program had a significant effect on postdoctoral teaching approaches and practices. There are inherent benefits to training graduate students and postdoctoral scholars before they enter academic careers (Austin 2011, Greer et al. 2016). The outcomes of FIRST IV have lasted well into participants’ faculty careers, an encouraging result for teacher professional development programs and lends support for the FIRST IV model of professional development.

In addition to providing evidence for professional development outcomes, our study uncovered important relationships among assessments of faculty and students. The results support the notion that individual’s teaching approaches are manifested in the classroom through their teaching practices. This informs future efforts to assess changes in teaching approach and possibly the effectiveness of teaching professional development. Learner-centered teaching practices were not only related to teaching approach, but were also detected by students across multiple faculty and institutions. These emergent relationships among assessments will prove valuable for future education research and the exploration of affecting change in instructor behavior, change, and student understanding in the classroom.

## Author contributions

NE collected data, analyzed the results, and wrote the manuscript.

JMM designed the study, collected data, and edited the manuscript.

DEM designed the study, collected data, and edited the manuscript.

## Acknowledgements

We thank the NSF for funding this study (DUE-1623834) and our advisory committee for helping steer the data collection and analysis of the results. We sincerely appreciate and thank all of the participating faculty and students who comprised the data collected in this study.

